# Identification of a Mitofusin specificity region that confers unique activities to Mfn1 and Mfn2

**DOI:** 10.1101/651851

**Authors:** SR Sloat, BN Whitley, EA Engelhart, S Hoppins

## Abstract

Mitochondrial structure can be maintained at steady state or modified in response to changes in cellular physiology. This is achieved by the coordinated regulation of dynamic properties including mitochondrial fusion, division and transport. Disease states, including neurodegeneration, are associated with defects in these processes. In vertebrates, two Mitofusin paralogs, Mfn1 and Mfn2, are required for efficient mitochondrial fusion. The Mitofusins share a high degree of homology and have very similar domain architecture, including an amino terminal GTPase domain and two extended helical bundles that are connected by flexible regions. Mfn1 and Mfn2 are non-redundant and are both required for mitochondrial outer membrane fusion. However, the molecular features that make these proteins functionally distinct are poorly defined. By engineering chimeric proteins composed of Mfn1 and Mfn2, we discovered a region that contributes to isoform-specific function (**Mitofusin I**soform **S**pecific **R**egion – MISR). MISR confers unique fusion activity and Mitofusin specific nucleotide-dependent assembly properties. We propose that MISR functions in higher order oligomerization either directly, as an interaction interface, or indirectly through conformational changes.

## INTRODUCTION

Membrane fusion is an essential process mediated by diverse proteins in eukaryotic cells. Initiation of membrane fusion usually involves a tethering event, which establishes a physical interaction between the fusion partners. To couple the tethered state to bilayer mixing, significant energetic barriers are overcome by the fusion machinery, often through large conformational changes. Although the final lipid mixing step is not well defined for any cellular membrane fusion event, dehydration and local destabilization of the bilayer by the fusion machinery is likely to contribute.

Mitochondrial fusion, division and transport are dynamic properties that coordinately maintain or modify the structure of mitochondria in cells (Chan, 2012; Friedman and Nunnari, 2014). The dynamic structure of mitochondria contributes to mitochondrial function and integrates into important cellular physiology, including cell cycle progression and cell death (Burté et al., 2015; Dorn, 2018; Pernas and Scorrano, 2016). The range of human diseases that result from mutations in the genes encoding the protein components of the fusion and division machinery highlights the importance of these processes (Itoh et al., 2013; Pareyson et al., 2015). Members of the dynamin-related protein family (DRP) mediate mitochondrial division as well as outer and inner membrane fusion. This diverse family of large GTPases remodel membranes by coupling stages of GTP hydrolysis to self-assembly and conformational changes (Antonny et al., 2016). For example, the mitochondrial division machine, Drp1, is first recruited to the mitochondrial surface by resident adaptor proteins. Once at the mitochondrial outer membrane, nucleotide-bound Drp1 assembles into a macromolecular structure that encircles the organelle and subsequent GTP hydrolysis triggers constriction and disassembly. In vertebrates, mitochondrial outer membrane fusion requires both Mitofusin1 and Mitofusin2 (Mfn1 and Mfn2), and inner membrane fusion is mediated by the protein Optic Atrophy 1 (Opa1). Interestingly, the presence of two outer membrane fusion DRPs is unique to vertebrate cells, as most model systems rely on a single Mitofusin, such as Fzo1 in *S. cerevisiae* or MARF in *D. melanogaster*.

As members of the DRP family, Mitofusins possess a large N-terminal GTPase domain. The GTPase catalytic cycle is expected to control Mitofusin activity through self-assembly and conformational changes that together drive membrane remodeling (Daumke and Roux, 2017). Although not highly similar to the Mitofusins, Atlastin is a DRP that mediates homotypic fusion of the endoplasmic reticulum. Atlastin tethers the two membranes of the fusion partners via an intermolecular GTPase domain interface and subsequent conformational changes induce lipid mixing, facilitated by the amphipathic helix on the C-terminus of Atlastin (McNew et al., 2013). The intermolecular GTPase domain interface represents a conserved feature of the DRP family and is required for nucleotide hydrolysis. This interface has been proposed to mediate Mitofusin-mediated membrane tethering (Cao et al., 2017; Yan et al., 2018), analogous to the Atlastin mechanism; however, a competing model implicates the C-terminal domain in this step (Franco et al., 2016; Koshiba et al., 2004).

Outside of the GTPase domain, relatively few functional domains have been identified and characterized in the Mitofusin proteins. The rest of the protein is predicted to be primarily alpha helical, with two predicted heptad repeat domains (HR1 and HR2) and a transmembrane region (Figure 1A). Although we lack complete structural information for the Mitofusins, recent atomic resolution structures of internally modified Mfn1 constructs reveal a globular GTPase domain connected to an extended helical bundle (HB1) composed of three helices from the first half of the protein and a single helix from the C-terminus (Cao et al., 2017; Qi et al., 2016; Yan et al., 2018). Based on homology to the bacterial dynamin-like protein (BDLP), the region absent from the structure is predicted to form a second extended helical bundle (HB2) (Low et al., 2009). The C-terminal domain of Mitofusin has been characterized to a greater extent, and is critical for function, but consensus on the topology of the protein is lacking. The structural data require that the C-terminal domain be cytoplasmic, which would result from a hairpin transmembrane anchor; however, recent biochemical analysis instead suggest that the membrane anchor is a single pass transmembrane helix, with the C-terminal domain residing in the intermembrane space (Mattie et al., 2018).

**Figure 1.**
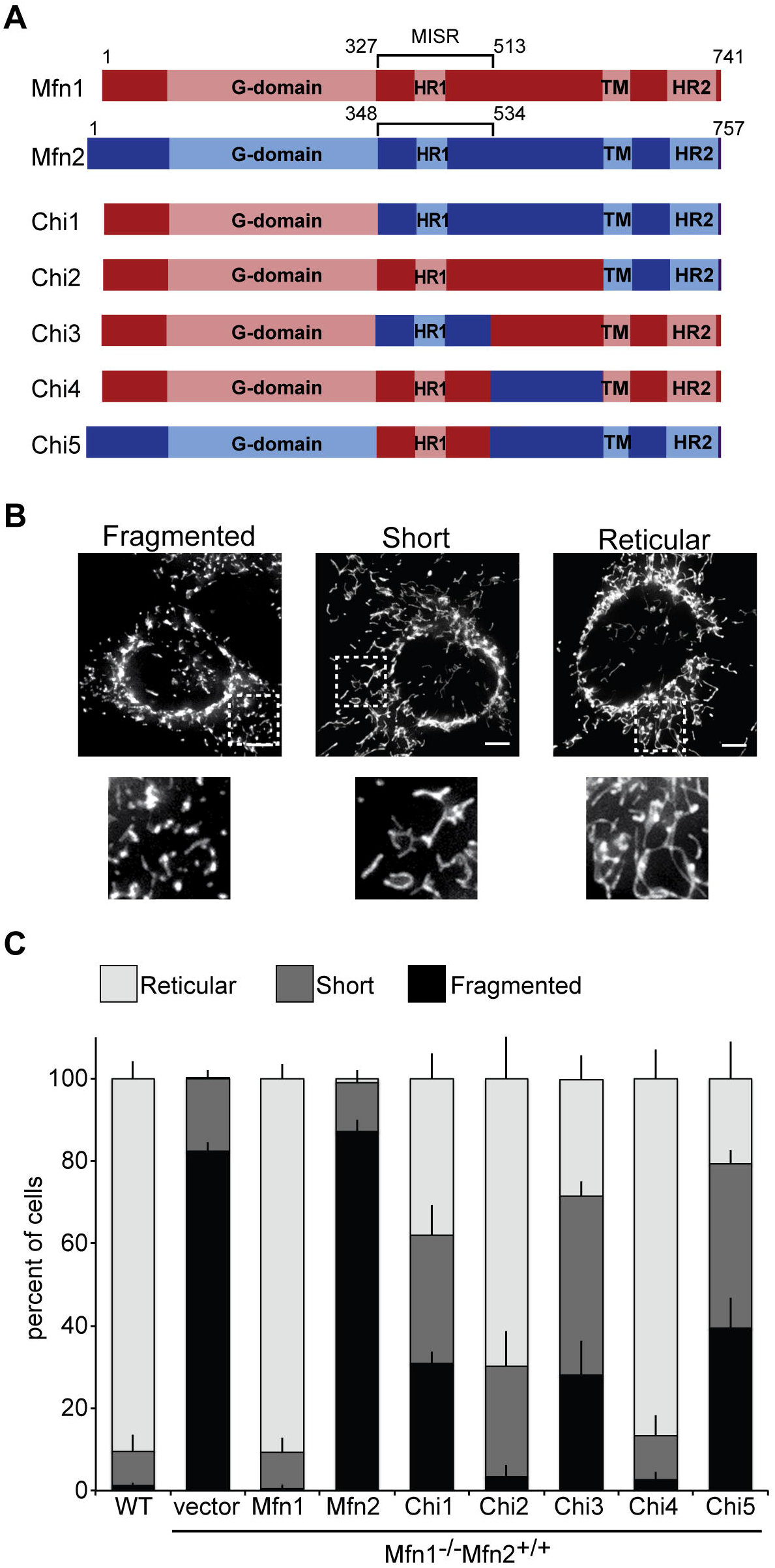
Mfn1-dependent rescue of Mfn1-null cells by Chimera proteins. (A) Schematic representation of known functional domains in Mfn1 and Mfn2 and the chimeric proteins generated for this study. (B) Representative images of MEF cells with mitochondrial structures that are scored as fragmented, short or reticular. Mitochondria were stained with Mitotracker Red CMXRos and visualized by fluorescence microscopy. Images represent a single plane from a Z-series. Scale bars are 5 μM. (C) Quantification of the mitochondrial morphology in cells from a clonal population of Mfn1-null cells expressing the indicated Mfn or Chi protein. Error bars indicate mean + standard deviation from three blinded experiments (n ≥ 100 cells per population per experiment).

While Mfn1 and Mfn2 share high primary sequence homology, they are expressed in a tissue specific manner (Santel and Fuller, 2001) and are functionally unique. This is underscored by the relatively low fusion efficiency of mitochondria that possess solely Mfn1 or Mfn2 (Chen et al., 2003; Chen et al., 2005). Indeed, Mfn1 and Mfn2 have been reported to possess unique GTPase and membrane tethering activities (Ishihara et al., 2004). Interestingly, both proteins are not required on both membranes of the fusion pair; rather, fusion efficiency is restored when Mfn1 and Mfn2 are present in trans, on opposite membranes of the fusion pair (Detmer and Chan, 2007; Hoppins et al., 2011). Mutations in Mfn2 are associated with the neurodegenerative disorder Charcot Marie Tooth syndrome Type 2A (CMT2A). Functional characterization of a subset of Mfn2 CMT2A variants revealed that several were unable to support fusion alone (Detmer and Chan, 2007). Surprisingly, the same CMT2A variants could support fusion activity when partnered with wild type Mfn1 in cis, on the same membrane, or in trans, on opposite membranes. Together, these data indicate that Mfn1 and Mfn2 contribute distinct molecular functions to the fusion complex. However, the biochemical properties of Mfn1 and Mfn2 remain poorly defined. To identify a domain that confers specific Mfn1 or Mfn2 activity, we generated chimeric proteins and characterized their function in cells and biochemically. Our analyses reveal a region that connects HB1 and HB2 and confers unique fusion activity to Mfn1 and Mfn2, likely through its role in nucleotide dependent self-assembly of each Mitofusin.

## RESULTS

### Identification of a region that confers Mitofusin specific function

The Mitofusin proteins are approximately 82% similar and 66% identical and have the same domain architecture. Therefore, we reasoned that the Mitofusin proteins would be amenable to domain swap experiments. To identify a domain that might confer Mitofusin-specific function, we generated chimeric proteins with regions of murine Mfn1 replaced with the equivalent segment from murine Mfn2 (Figure 1A). To functionally assess these chimeric proteins, each was expressed in mouse embryonic fibroblast (MEF) cell lines that lack Mfn1 (Chen et al., 2003). Loss of Mfn1 disrupts the steady-state reticular mitochondrial network due to low levels of mitochondrial fusion and ongoing division resulting in many small mitochondrial fragments. It has been previously demonstrated that, even when mildly overexpressed, Mfn2 cannot fully restore mitochondrial fusion activity in Mfn1-null cells (Chen et al., 2003). We therefore reasoned that the exchange of a region that confers Mitofusin specific function would result in a chimeric protein with unique fusion activity that could be detected in the Mfn1-null cells.

We first sought to establish a system to test the activity of our chimeric proteins. As overexpression of Mitofusin can also alter the function of the protein (Eura et al., 2003; Rojo et al., 2002), we analyzed cells expressing Mitofusin proteins at near-endogenous levels. We stably expressed Mitofusin chimeric proteins with a C-terminal FLAG tag in Mfn1-null cells by retroviral transduction and screened clonal populations for those with Mitofusin expression levels comparable to wild type for further analysis (Supplemental Figure S1). These stable cell lines were examined in blinded experiments to score the mitochondrial morphology in each cell as either as reticular, short or fragmented (Figure 1B)(reticular mitochondria are > 7.5 microns; short mitochondria are between 2.5 and 7.5 microns; fragmented mitochondria are smaller than 2.5 microns). The mitochondrial network of wild type MEF cells is reticular with very few short or fragmented organelles observed and mitochondria that are highly connected (Figure 1C). The mitochondria in Mfn1-null cells transduced with empty vector were fragmented, as expected for cells expressing only Mfn2. In contrast, expression of Mfn1-FLAG restored a connected reticular network in the vast majority of cells, so that these stable lines were indistinguishable from wild type controls (Figure 1C and Supplemental Figure S1). However, expression of Mfn2-FLAG in Mfn1-null cells did not alter the mitochondrial structure significantly, as these cells were comparable to vector controls (Figure 1C and Supplemental Figure S1). Together, these data are consistent with previously reported observations (Chen et al., 2003) and established the basis for our functional screen.

In the first pair of chimeric proteins, we replaced either the entire protein excluding the GTPase domain or the transmembrane domain and the C-terminus of Mfn1 with the equivalent portion of Mfn2 to generate Chimera1 and Chimera2 (Figure 1A, Chi1 and Chi2, respectively). The chimeric proteins were expressed in Mfn1-null cells (with endogenous Mfn2) and clonal populations with expression comparable to wild type were chosen for further analysis (Supplemental Figure S1). We observed that more than half of Mfn1-null cells expressing Chi1 possessed mitochondria that were either fragmented or short (Figure 1C). This indicates that Chi1 did not support robust fusion in Mfn1-null cells. In contrast, the majority of cells expressing Chi2 were found to have a reticular mitochondrial network (Figure 1C). Thus, Chi2 was sufficient to restore fusion activity. We obtained similar results with two additional clonal populations for each construct (Supplemental Figure S2).

Together, the different functional activity of Chi1 and Chi2 suggest that the C-terminal domain of Mfn1 and Mfn2 is not functionally unique, while the region between the GTPase domain and the transmembrane region does confer unique fusion activity. To further dissect the region, we analyzed the N- and C-terminal segments of this poorly characterized middle domain. We utilized a homology model of the Mitofusins based on full length BDLP (PDB 2J68) and identified a predicted unstructured loop that divided this domain into two pieces (Supplemental Figure S3). From this, we generated two chimeric proteins, Chimera3 and Chimera4, in which the N- and C-terminal halves of the middle domain of Mfn2 replace the equivalent pieces of Mfn1 (Chi3 and Chi4, respectively) (Figure 1A). As described above, we generated clonal populations of Mfn1-null cells expressing each chimera at levels comparable to endogenous Mfn1 (Supplemental Figure S1). Most cells expressing Chi3 had short or fragmented mitochondria (Figure 1C). In contrast, the expression of Chi4 resulted in many cells with reticular mitochondria, comparable to both Mfn1 and Chi2. Similar results were also observed in two additional clonal populations for each construct (Supplemental Figure S2). Together, these data suggest that Chi3 contains a Mitofusin isoform-specific region (MISR). Based on the homology model with BDLP and the truncated Mfn1 structure, this region contains part of a helix from HB1 and two helices connected by a loop within HB2 followed by another large loop (Supplemental Figures S3 and S4, highlighted in purple).

As an additional test of whether this domain confers Mitofusin-specific function, we set out to test if MISR was sufficient to alter the fusion activity of Mfn2 when expressed in Mfn1-null cells. We expected that the presence of a region with Mfn1-specific function would improve the fusion efficiency of the chimeric protein compared to Mfn2. To test this, we generated a chimeric protein in which Mfn2 MISR was replaced with Mfn1 MISR (Chimera5) (Figure 1A, Chi5). Compared to Mfn1-null cells expressing Mfn2-FLAG, which have fragmented mitochondria, cells expressing Chi5 more frequently had short or reticular mitochondria (Figure 1C). A similar pattern was observed in two additional clonal populations (Supplemental Figure S2). This indicates that Chi5 supports more fusion with endogenous Mfn2 in Mfn1-null cells than Mfn2-FLAG. Therefore, MISR confers a Mitofusin-specific function with unique mitochondrial fusion efficiency in Mfn1-null cells.

### Chimeric proteins possess unique fusion characteristics

To characterize Chi3 and Chi5 function in more detail, we quantified the mitochondrial fusion activity supported by the chimeric proteins utilizing an established *in vitro* mitochondrial fusion assay (Hoppins et al., 2011). This approach provides a quantitative measure of Mitofusin-mediated mitochondrial fusion in the absence of other contributing factors in cells such as microtubule dependent transport or cytosolic pro-fusion proteins, such as Bax (Hoppins et al., 2011; Karbowski et al., 2006). We utilized the same cell lines described in Figure 1. Mitochondria were isolated from cells stably expressing either mitochondrially targeted CFP (mtCFP) or TagRFP (mtRFP), mixed together in the presence of fusion buffer and analyzed by fluorescence microscopy following incubation. Fusion events were defined by colocalization of CFP and RFP in a single organelle, and fusion efficiency is expressed as a proportion of wild type controls performed in parallel. First, we assessed homotypic fusion efficiency, which reflects the Mitofusin composition of the cells scored in Figure 1. In these reactions, both cyan and red mitochondria possess the same Mitofusin proteins (Figure 2A, black bars). As expected, mitochondria isolated from Mfn1-null cells stably expressing Mfn1-FLAG have fusion activity comparable to wild type controls (Figure 2A, black bars, Mfn1*-Mfn2 and Mfn1-Mfn2, respectively). In contrast, mitochondria isolated from Mfn1-null cells transduced with either empty vector or Mfn2-FLAG fused poorly compared to wild type controls (Figure 2A, black bars, Mfn2 and Mfn2*-Mfn2, respectively). In both of these reactions only Mfn2 is present; therefore, these data are consistent with our previous observation that mitochondrial fusion is inefficient when Mfn2 is present but Mfn1 is not (Hoppins et al., 2011).

**Figure 2.**
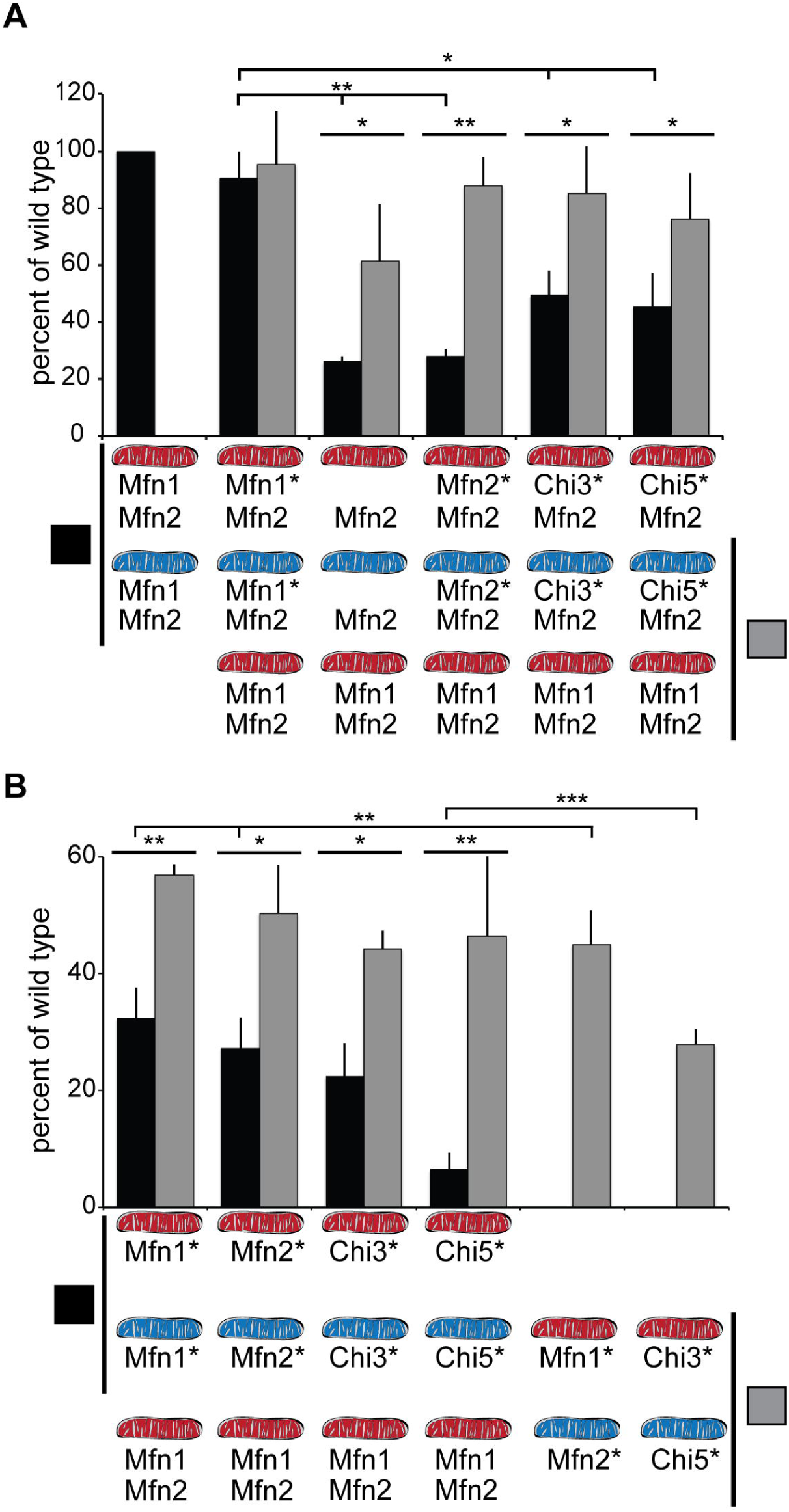
Fusion activity of Chi proteins in heterotypic and homotypic complexes as assessed in vitro. (A) Mitochondria were isolated from wild type cells (Mfn1 Mfn2) or clonal populations of Mfn1-null cells transduced with empty vector (Mfn2) or expressing either Mfn1-FLAG (Mfn1*Mfn2), Mfn2-FLAG (Mfn2*Mfn2), Chi3-FLAG (Chi3*Mfn2) or Chi5-FLAG (Chi5*Mfn2), where the asterisk indicates the non-endogenous protein. The indicated mitochondrial combinations were subject to in vitro fusion conditions at 37°C for 60 minutes and data is expressed as a relative amount of wild type controls, performed in parallel. Black bars indicate homotypic reactions, where both mitochondrial fusion partners possess the same Mitofusin proteins. Grey bars indicate heterotypic fusion reactions, where one of the mitochondrial fusion partners is wild type. Error bars indicate mean + standard deviation from at least four independent experiments and the statistical significance indicated on the graphs were determined by paired t-test analysis (one tail); *P<0.05, **P<0.005. (B) Mitochondria were isolated from wild type cells (Mfn1 Mfn2) or clonal populations of Mfn1/2-null cells expressing either Mfn1-FLAG (Mfn1*), Mfn2-FLAG (Mfn2*), Chi3-FLAG (Chi3*) or Chi5-FLAG (Chi5*), where the asterisk indicates the non-endogenous protein. The indicated mitochondrial combinations were subject to in vitro fusion conditions at 37°C for 60 minutes and data is expressed as a relative amount of wild type controls, performed in parallel. Black bars indicate homotypic reactions, where both mitochondrial fusion partners possess the same Mitofusin proteins. Grey bars indicate heterotypic fusion reactions. Error bars indicate mean + standard deviation from at least four independent experiments and the statistical significance indicated on the graphs were determined by paired t-test analysis (one tail); *P<0.05, **P<0.005, ***P<0.0005.

To test the fusion activity of Chi3 or Chi5, we isolated mitochondria from Mfn1-null cells expressing the chimeric protein with endogenous Mfn2. Mitochondria with Chi3 and Mfn2 had fusion activity that was lower than wild type controls, consistent with the partial rescue of mitochondrial morphology shown in Figure 1 (Figure 2A, black bars, Chi3*-Mfn2). Mitochondria with Chi5 and Mfn2 fused more than mitochondria with only Mfn2, but not as well as wild type controls (Figure 2A, black bars, Chi5*-Mfn2). The comparable fusion activity of Chi3-Mfn2 and Chi5-Mfn2 mitochondria is consistent with the similar mitochondrial morphology of these clonal populations presented in Figure 1. Together, these *in vitro* experiments support the conclusions from our *in vivo* assays, which indicate that MISR impacts the unique fusion activity of Mfn1 and Mfn2.

We have previously shown that the fusion efficiency of mitochondria that possess only Mfn1 or Mfn2 increased when a wild type fusion partner was provided (Hoppins et al., 2011). This is due to the presence of the opposite isoform on the wild type mitochondria, generating a heterotypic fusion complex. To determine if wild type mitochondria could improve the fusion efficiency of the Chi3/5-Mfn2 mitochondria in trans, we performed another set of *in vitro* fusion reactions that paired mitochondria isolated from the clonal populations of Mitofusins in Mfn1-null cells with wild type mitochondria (Figure 2A, grey bars). As expected, the fusion efficiency of Mfn1-FLAG-Mfn2 mitochondria did not change with a wild type partner (Figure 2A, grey bar, Mfn1*-Mfn2). Consistent with previously published data, the fusion efficiency of mitochondria bearing only Mfn2 was significantly increased by a wild type fusion partner (Figure 2A, grey bars, Mfn2 and Mfn2*-Mfn2). The relative amount of fusion also increased for Chi3-Mfn2 mitochondria and Chi5-Mfn2 mitochondria when a wild type partner was provided in trans (Figure 2A, grey bars, Chi3*-Mfn2 and Chi5*-Mfn2).

These experiments suggest that, similar to Mfn1 and Mfn2, the chimeric proteins support the most fusion in the context of a heterotypic fusion complex. Next, we wanted to examine the fusion characteristics of both Chi3 and Chi5 alone. Cells expressing only one Mitofusin or chimeric protein were created by retroviral transduction of Mfn1-FLAG, Mfn2-FLAG, Chi3-FLAG or Chi5-FLAG into Mfn1^−/−^Mfn2^−/−^ cells (Mfn1/2-null). As described above for the Mfn1-null cells, clonal populations of stable cell lines were screened for expression of the Mitofusin or chimera by western blot analysis of whole cell extracts, and isolates with expression levels near wild type were chosen for further analysis (Supplemental Figure S5A). These stable cell lines were examined in blinded experiments to quantify mitochondrial morphology. As expected, almost all wild type cells had a connected, reticular mitochondrial network and all Mfn1/2-null cells transduced with empty vector had a fragmented mitochondrial network (Supplemental Figure S5A). Approximately half of the cells expressing either Mfn1 or Mfn2 at near endogenous levels had reticular or short mitochondria, consistent with low levels of fusion activity by homotypic complexes (Supplemental Figure S5B). Others have reported that expression of either Mfn1 or Mfn2 can restore a reticular network in Mfn1/2-null cells; we suggest that the difference is due to lower expression of the Mitofusin protein in our approach (Cao et al., 2017) (Yan et al., 2018). In contrast to Mfn1 and Mfn2, only 25% of cells expressing Chi3 at near endogenous levels possessed reticular or short mitochondria and the vast majority of cells expressing Chi5 had a fragmented mitochondrial network (Supplemental Figure S5B).

These clonal cell lines expressing a single Mitofusin were then utilized to quantify the mitochondrial fusion efficiency *in vitro*. First we assessed homotypic fusion reactions, with the same Mitofusin protein present on all mitochondria in the fusion reaction (Figure 2B, black bars). Mitochondria expressing only Mfn1-FLAG or Mfn2-FLAG support less fusion than wild type control mitochondria (Figure 2B, Mfn1, black bar). Mitochondria with only Chi3 fused as well as the Mfn1-FLAG and Mfn2-FLAG controls, demonstrating that this chimeric protein has fusion activity similar to Mfn1 or Mfn2 in the context a homotypic complex (Figure 2B, Chi3, black bar). In contrast, mitochondria with only Chi5 fused poorly, indicating that this chimeric protein supports very limited mitochondrial fusion in the absence of another Mitofusin (Figure 2B, Chi5, black bar).

To determine if, like Mfn1 and Mfn2, Chi3- or Chi5-mediated fusion is increased when Mfn1 and Mfn2 are present in trans on the fusion partner, we performed heterotypic *in vitro* fusion reactions with wild type mitochondria. As expected, if we pair mitochondria that have only Mfn1-FLAG or Mfn2-FLAG with wild type mitochondria, we observed higher rates of fusion than in homotypic reactions (Figure 2B, Mfn1*, Mfn2*; compare grey and black bars, respectively). Similarly, mitochondria expressing only Chi3 or Chi5 fused more with a wild type partner (Figure 2B Chi3* or Chi5*, compare grey bars [heterotypic] and black bars [homotypic]). Interestingly, despite the poor fusion of mitochondria with Chi5 alone, fusion efficiency with a wild type partner was similar to Mfn1 and Chi3, indicating that Chi5 has comparable fusion activity only when another Mitofusin is present, either on the same membrane or on opposite membranes.

We have previously reported that by combining mitochondria that possess only Mfn1 with mitochondria that possess only Mfn2, *in vitro* mitochondrial fusion activity is significantly higher than homotypic reactions (Hoppins et al., 2011). We have recapitulated that effect here with mitochondria isolated from the clonal populations of Mfn1/2-null cells expressing either Mfn1 or Mfn2 at near endogenous levels (Figure 2B). To determine if Chi3 and Chi5 also work more effectively as a heterotypic complex in trans, we quantified in vitro mitochondrial fusion with this pairing. The fusion efficiency of Chi5 is significantly higher when Chi3 is present on the fusion partner, compared to homotypic Chi5 reactions. Therefore, by assessing the fusion activity of mitochondria containing a single Mitofusin protein, our data reveal that Chi3 and Chi5 possess unique fusion properties, which is consistent with MISR conferring unique function to the Mitofusin proteins.

### In a heterotypic complex with Mfn1, membrane fusion by Chi5 requires a functional GTPase domain while Mfn2 does not

The unique molecular contributions of Mfn1 and Mfn2 to the heterotypic fusion complex are further highlighted by the characterization of mutant variants of Mfn2 associated with CMT2A. Specifically, some mutant variants that restored a reticular network when slightly overexpressed in Mfn2-null cells had no fusion activity in Mfn1/2-null cells (Detmer and Chan, 2007). These data indicate that the disease-associated variants contributed to fusion activity with endogenous Mfn1 protein in the Mfn2-null cells, but the same mutant variant could not mediate fusion alone in Mfn1/2-null cells. We set out to determine if this Mfn2-specific fusion activity was retained in Chi5.

As positive controls, we recapitulated published data by expressing mutant variants of Mfn2 with a C-terminal mNeonGreen tag (Mfn2-NG) in Mfn2-null cells and scored mitochondrial morphology in blinded experiments. As expected, Mfn2-null cells transduced with empty vector predominantly possessed mitochondria that were fragmented (Figure 3A). In contrast, the majority of cells expressing Mfn2-NG had mitochondrial networks that were highly connected and reticular, comparable to wild type controls. The first mutant variant that we assessed was Mfn2-K109A, which is predicted to bind, but not hydrolyze GTP (Yan et al., 2018). In just under half of the cells expressing Mfn2-NG-K109A, the mitochondrial network remained highly fragmented, similar to previously published results (Detmer and Chan, 2007). Next, we expressed Mfn2-R94W, which is the most frequently observed position affected in CMT2A patients. Also consistent with previously published data, expression of Mfn2-NG-R94W in Mfn2-null cells restored a reticular mitochondrial network in the vast majority of cells (Figure 3A).

**Figure 3.**
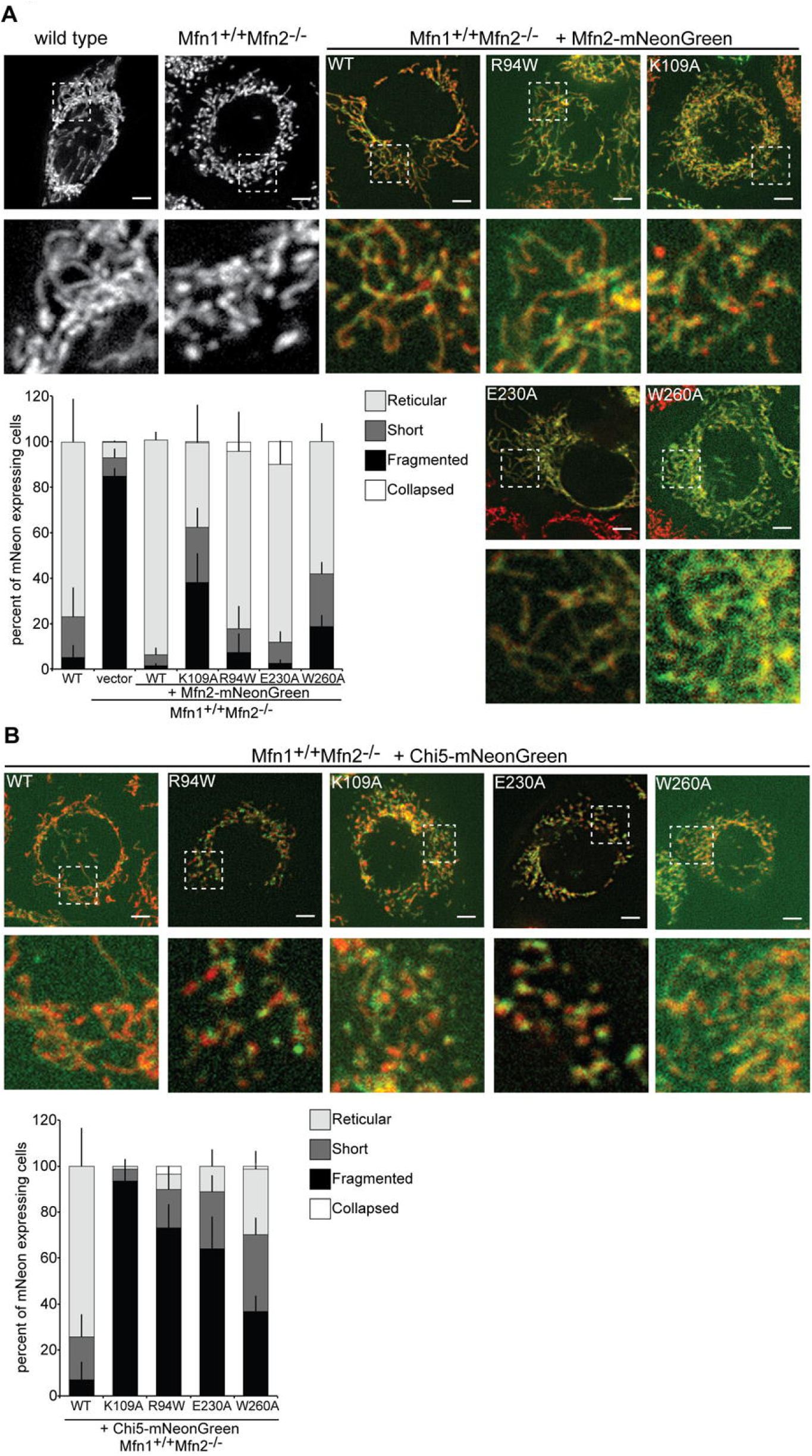
Mfn2 GTPase mutant variants support fusion with endogenous Mfn1, but Chi5 variants do not. (A) Representative images of wild type and Mfn1^+/+^ Mfn2^−/−^ cells transduced with either empty vector or Mfn2-NG. Mitochondria were labeled with MitoTracker Red CMXRos and visualized by fluorescence microscopy. Images represent maximum intensity projections. Scale bars are 5 µM. Quantification of the mitochondrial morphology is shown in the graph. Error bars indicate mean + standard deviation from three blinded experiments (n ≥ 100 cells per population per experiment). (B) Representative images of Mfn1^+/+^Mfn2^−/−^ cells transduced with Chi5-NG. Mitochondria were labeled with MitoTracker Red CMXRos and visualized by fluorescence microscopy. Images represent maximum intensity projections. Scale bars are 5 µM. Quantification of the mitochondrial morphology is shown in the graph. Error bars indicate mean + standard deviation from three blinded experiments (n ≥ 100 cells per population per experiment).

To further test for the requirement of GTPase activity in Mfn2 function, we examined the fusion activity of mutant variants of Mfn2 with substitutions of key catalytic residues identified in the partial structure of Mfn1. We constructed Mfn2 mutant variants with substitutions in two residues critical for GTP hydrolysis in Mfn1: Mfn2-E230A (Mfn1-E209A) and Mfn2-W260A (Mfn1-W239A)(Cao et al., 2017). Based on the structure, E230/E209 contributes to the intermolecular interface of the GTPase domain. The W260/W239 side chain has been reported to be located in the nucleotide binding pocket or at the GTPase interface (Cao et al., 2017; Yan et al., 2018). Importantly, when these mutant variants were expressed in the context of full length Mfn1 or Mfn2 protein in Mfn1/2-null cells, none of them supported fusion, consistent with loss of enzyme activity (Cao et al., 2017; Yan et al., 2018). Here, we expressed either Mfn2-NG-E230A or Mfn2-NG-W260A in Mfn2-null cells. In both cases, the mutant variants robustly supported mitochondrial fusion with endogenous Mfn1, as we observed that most cells expressing the variant had a connected mitochondrial network (Figure 3A). Taken together, these data support the conclusion that a mutant variant of Mfn2 that lacks normal catalytic activity can contribute to fusion activity when partnered with wild type Mfn1.

To determine if Chi5 GTPase mutant variants can mediate fusion with endogenous Mfn1, we generated the same four mutants in Chi5 and expressed these as C-terminal mNeonGreen fusion proteins (Chi5-NG) in Mfn2-null cells (Figure 3B). The expression of wild type Chi5-NG in Mfn2-null cells generated a reticular, connected mitochondrial network in most cells, indicating that Chi5 supports robust fusion with endogenous Mfn1 (Figure 3B). In contrast, the mitochondrial network remained fragmented in cells expressing Chi5-NG-K109A, Chi5-NG-R94W, Chi5-NG-E230A or Chi5-NG-W260A. These data indicate that, unlike Mfn2, the mutant Chi5 variants cannot mediate fusion with a wild type Mfn1 partner. This is consistent with our conclusion that Chi5 is functionally distinct from Mfn2 and that MISR is important to confer Mitofusin-specific functional activity.

### Nucleotide dependent assembly is a unique property of Mfn1 and Mfn2

Mfn1 and Mfn2 have been reported to physically interact (Detmer and Chan, 2007; Ishihara et al., 2004; Qi et al., 2016). To determine if Chi3 and Chi5 interact with endogenous Mfn2 in Mfn1-null cells, we performed co-immunoprecipitation from mitochondria isolated from the clonal populations. As expected, both Mfn1-FLAG and Mfn2-FLAG pull down endogenous Mfn2, but not VDAC, another abundant mitochondrial outer membrane protein (Supplemental Figure S6). Furthermore, Chi3 and Chi5 also interact with Mfn2, consistent with our cellular and in vitro analysis of mitochondrial fusion indicating that these proteins are functional Mitofusin variants.

One important aspect of DRP function in membrane remodeling is nucleotide-dependent self-assembly into higher order structures. To date, the only domain experimentally demonstrated to be associated with assembly of the Mitofusins is the GTPase domain, which forms an intermolecular dimer required for nucleotide hydrolysis (Cao et al., 2017; Qi et al., 2016; Yan et al., 2018). We therefore assayed nucleotide dependent assembly of wild type and chimeric Mitofusin proteins utilizing blue native polyacrylamide gel electrophoresis (BN-PAGE), which has previously detected Mitofusin complexes ranging from ∼140 – 440 kilodaltons (kDa) (Ishihara et al., 2004; Karbowski et al., 2006; Steffen et al., 2017).

To evaluate the assembly state of Mfn1, Mfn2, Chi3 and Chi5 in the mitochondrial outer membrane, mitochondria were isolated from the Mfn1-null clonal populations that were utilized for the in vitro fusion analysis in Figure 2A. These mitochondria were either left untreated or were incubated with the indicated nucleotide conditions before detergent solubilization and BN-PAGE followed by western blot analysis. In the untreated samples, the majority of Mfn1-FLAG and Mfn2-FLAG were detected in a single oligomeric state that we predict to be a dimer that migrates as a ∼200 kDa species (Figure 4A, line). The addition of GDP alone did not alter the assembly state of the Mitofusins (Figure 4A). In contrast, addition GDP and BeF_3_, which mimics the transition state when GDP and the free gamma phosphate are still both in the nucleotide-binding pocket, caused Mfn1-FLAG to shift dramatically from the dimer into two higher molecular weight oligomeric species (Figure 4A, ∼320 kDa [double arrow] and ∼450 kDa [arrow]). Although Mfn2-FLAG was also detected in higher order oligomers in the presence of the transition state mimic, there were notable differences from Mfn1 assembly. Specifically, Mfn2 did not form significant quantities of the 450 kDa species that is formed by Mfn1 under these conditions (Figure 4A, compare arrow). Together, these data indicate that the Mitofusins undergo nucleotide dependent assembly, and that Mfn1 and Mfn2 respond differentially to the same nucleotide state. Thus, nucleotide-dependent assembly is a Mitofusin-specific function where Mfn1 robustly forms three oligomeric species and Mfn2 predominantly forms two.

**Figure 4.**
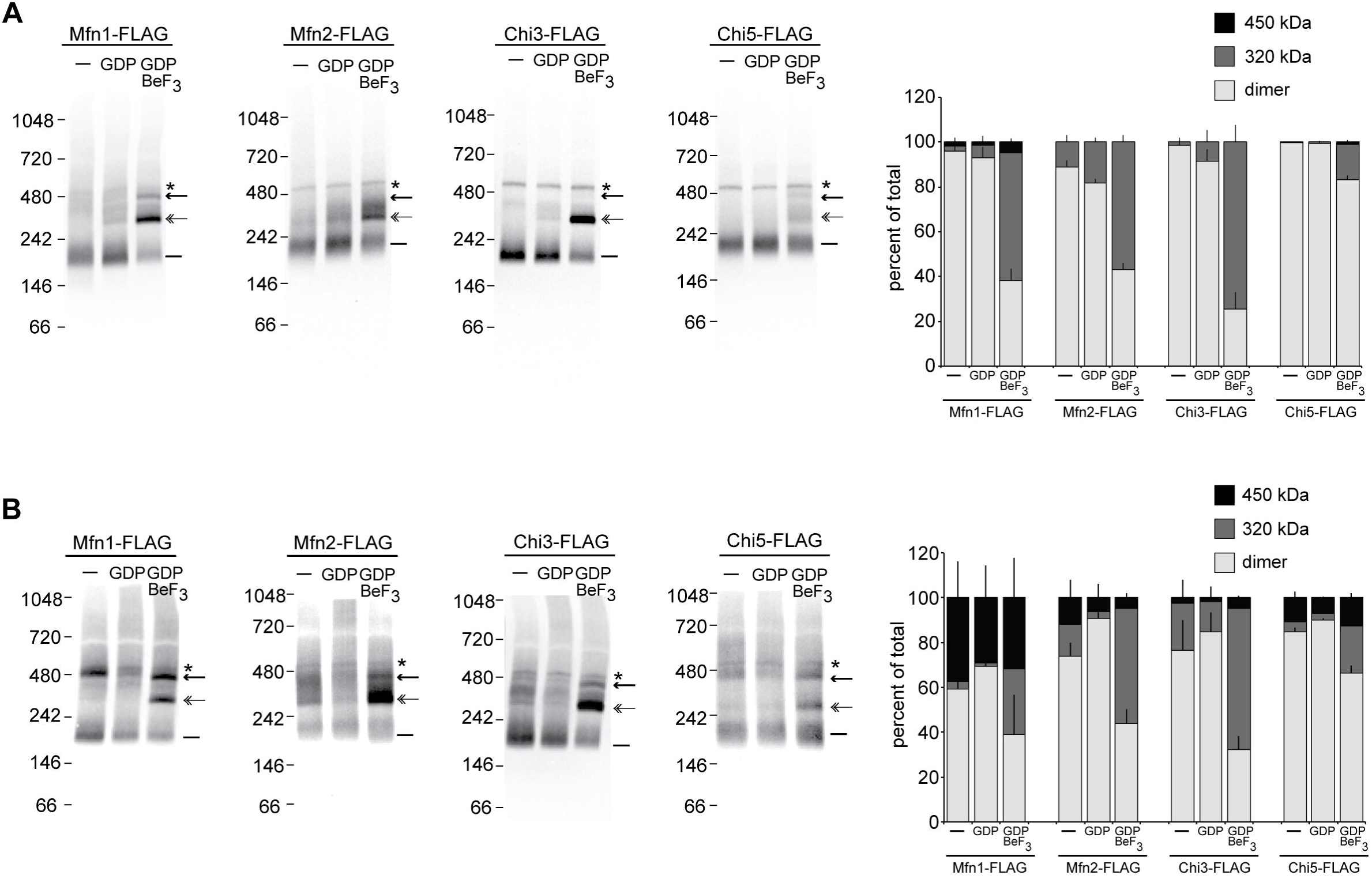
Mitofusin-specific nucleotide dependent assembly. (A) Mitochondria were isolated from clonal populations of Mfn1-null cells expressing Mfn1-FLAG, Mfn2-FLAG, Chi3-FLAG or Chi5-FLAG. Mitochondria were either untreated (−) or incubated with the specified nucleotide conditions and subsequently subjected to detergent solubilization and analysis by BN-PAGE and immunoblotting with anti-FLAG. The positions of the molecular weight markers are shown on the left. The predicted dimer is indicated with a line ^∼^(); the ∼320 kDa species is indicated with a double arrow; the ∼450 kDa species is indicated with an arrow; a non-specific band is highlighted with an asterisk (*). The percentage of total protein in each oligomeric state for each condition is represented in the bar graph as mean + standard deviation of three independent experiments. (B) Mitochondria were isolated from clonal populations of Mfn1/2-null cells expressing Mfn1-FLAG, Mfn2-FLAG, Chi3-FLAG or Chi5-FLAG. Mitochondria were either untreated (−) or incubated with the specified nucleotide conditions and subsequently subjected to detergent solubilization and analysis by BN-PAGE and immunoblotting with anti-FLAG. The positions of the molecular weight markers are shown on the left. The predicted dimer is indicated with a line ^∼^(); the ∼320 kDa species is indicated with a double arrow; the ∼450 kDa species is indicated with an arrow; a non-specific band is highlighted with an asterisk (*). The percentage of total protein in each oligomeric state for each condition is represented in the bar graph as mean + standard deviation of three independent experiments.

As expected, both Chi3-FLAG and Chi5-FLAG migrated primarily as dimers in untreated and GDP samples (Figure 4A, line). In the presence of the transition state mimic, Chi3 assembled robustly into the 320 kDa species. However, unlike Mfn1, Chi3 does not readily form the 450 kDa species and in this way, is similar to Mfn2 (Figure 4A, arrow). The Chi5 protein showed the lowest propensity to assembly in the presence of the transition state mimic, with most of the Chi5 protein in the dimer species under all conditions (Figure 4A and 4B, line). Interestingly, Chi5 forms some 450 kDa species, suggesting that Mfn1 MISR in Chi5 supports this assembly state (Figure 4A, arrow). Given the low propensity of Chi5 to assemble, we assessed its topology in the mitochondrial outer membrane by protease protection. Chi5 is targeted to mitochondria, as the protein abundance in untreated mitochondria was comparable to Mfn2 controls (Supplemental Figure S7A). As observed with the mitochondrial outer membrane protein Tom20, both Mfn2-FLAG and Chi5-FLAG are completely digested by the addition of proteinase K (PK) to intact mitochondria, indicating that both have the same topology. These data are not consistent with recent reports that indicate that the C-terminus of Mitofusins are localized to the intermembrane space (Mattie et al., 2018), but are consistent with structural data where the C-terminus of Mitofusin is a component of HB1 in the cytosol (Cao et al., 2017; Yan et al., 2018). Therefore, despite its correct targeting and assembly in the mitochondrial outer membrane, Chi5 has limited nucleotide dependent assembly, which is likely due to MISR, as this is the only discernable difference with Mfn2. This inefficient assembly of Chi5 may be responsible, at least in part, for the poor fusion activity of Chi5 homotypic complexes reported in Figure 2B.

To determine if the nucleotide-dependent assembly of Mitofusin observed here by BN-PAGE requires both isoforms, we assessed the assembly in mitochondria that express a single Mitofusin protein. To do this, mitochondria were isolated from the Mfn1/2-null clonal populations that were utilized for the *in vitro* fusion analysis in Figure 2B. In the untreated samples, most Mfn1 migrated as a dimer, but some protein was detected in the two higher order species (Figure 4B, line and arrow). This pattern did not change significantly following incubation with GDP. With the addition of the transition state mimic, Mfn1 is distributed between all three oligomeric species (Figure 4B, Mfn1-FLAG). Although the relative abundance of the different Mfn1 oligomers was slightly different compared to those in Figure 4A, the size of each was indistinguishable (compare Figure 4B and 4A, respectively). For Mfn2, in either untreated or GDP-treated mitochondria, the majority of the protein migrated as a predicted dimer, with some in higher order assemblies (Figure 4B, Mfn2-FLAG). In contrast, most of the protein migrated as predicted trimer in the presence of the transition state mimic, consistent with the pattern observed with Mfn1-null mitochondria (compare Figure 4B and 4A, respectively). Consistent with our previous report, these data indicate that these oligomers contain either Mfn1 or Mfn2, but not both (Engelhart and Hoppins, 2019). Interestingly, for both Mfn1 and Mfn2, we observe more of the 450 kDa species when only a single Mitofusin is expressed compared to conditions where both Mitofusin proteins are expressed.

In the untreated and GDP-treated samples of mitochondria isolated from Mfn1/2-null expressing Chi3, the protein migrates primarily as a dimer and shifts into the 320 kDa species following incubation with the transition state mimic (Figure 4B, Chi3-FLAG). Despite being primarily Mfn1 amino acid composition, the assembly of Chi3 is more similar to Mfn2, which also assembles primarily into the 320 kDa species when incubated with the transition state mimic and has minor quantity of the 450 kDa species. The nucleotide-dependent assembly of Chi5 is more easily detected when this protein is expressed alone in Mfn1/2-null cells (Figure 4B, Chi5-FLAG). In contrast to the dimer observed with untreated and GDP-treated mitochondria, incubation with the transition state mimic resulted in Chi5 forming similar amounts of the 320 kDa and 450 kDa oligomers. This assembly pattern more closely resembles Mfn1 than Mfn2, despite Chi5 being primarily Mfn2 amino acid sequence. Together, this suggests that MISR plays a role in nucleotide dependent assembly. We propose that Mfn1-MISR promotes the assembly of the 450 kDa species, which is more abundant in Mfn1 than Mfn2. This conclusion is supported by the chimeric protein analysis as Chi5 forms more of the 450 kDa species than Mfn2 and Chi3 forms less of the 450 kDa species is formed than Mfn1 under the same conditions.

## DISCUSSION

By assessing the function of chimeric proteins composed of both Mfn1 and Mfn2, we have identified a region that confers a Mitofusin isoform-specific function. Exchanging this domain between Mfn1 and Mfn2 generated chimeric proteins with unique fusion activity and nucleotide dependent self-assembly characteristics. Within the linear protein sequence, this region is adjacent to the GTPase domain including most of alpha helix 3; however, the bulk of this region is within the poorly characterized predicted HB2 structure. In other DRP proteins, such as dynamin itself, multiple protein-protein interaction surfaces are found in the extended helical bundle. Therefore, MISR could contribute to an assembly surface in the Mitofusin proteins. Our data indicate that these are not Mfn1-Mfn2 heterotypic complexes, but we cannot rule out the possibility that another unknown protein is a component of one or more of the oligomers. Alternatively, in the related BDLP from cyanobacteria, the connection between HB1 and HB2 represents a functionally significant hinge that mediates a switch from open to closed conformations (Low et al., 2009). Given that MISR includes the loop and helices that compose part of the hinge, MISR could impact conformational changes in full length Mitofusin and thus influence nucleotide-dependent assembly via Mitofusin specific conformational flexibility. Given the drastic differences in these conformational states, it is possible that the 320 kDa and 450 kDa oligomers observed by BN-PAGE represent a tetramer in different conformational states.

Together with previously published data, our analyses indicate that GTPase deficient Mfn2 can contribute to fusion when paired with Mfn1. This is similar to the yeast inner membrane fusion machine, Mgm1, which only requires GTPase activity in the short, soluble form for membrane fusion to occur in cells (DeVay et al., 2009). In contrast, when the same mutants are expressed in Mfn2-null cells in the context of Chi5, the mutant variant did not mediate fusion, despite being primarily Mfn2 amino acid composition. Therefore, the Mfn1-specific molecular activity conferred MISR makes the GTPase mutations null alleles. We predict that the unique conformational flexibility and/or assembly interfaces contributed by MISR underlie this behavior.

Fusion is most effective when both Mfn1 and Mfn2 are present, likely due to their unique molecular functions. We discovered MISR by assessing the function of the chimera proteins in the presence of endogenous Mfn2, in Mfn1-null cells. As is the case for Mfn1 and Mfn2, both Chi3 and Chi5 fuse most effectively when another Mitofusin is present. Interestingly, these chimeric proteins have distinct characteristics in homotypic fusion reactions, where the same single Mitofusin is on both mitochondria in the fusion pair. Mitochondria with Chi3 had relative fusion that was comparable to mitochondria with only Mfn1, while mitochondria with only Chi5 fused very infrequently. Given that Chi5 also demonstrated less efficient nucleotide-dependent assembly, we predict that the assembly from dimers to higher-order species plays an important role in Mitofusin-mediated membrane fusion. Interestingly, the impact of less efficient assembly on mitochondrial fusion efficiency is less apparent in mitochondria with both Chi5 and Mfn2. This further highlights that in vertebrates, mitochondrial fusion efficiency depends on the expression of both functionally distinct Mitofusin isoforms.

### Abbreviations

Mfn Mitofusin; MISR Mitofusin isoform specific region; DRP dynamin-related protein; CMT2A Charcot Marie Tooth Syndrome Type 2A; HB helical bundle; MEF mouse embryonic fibroblast; BN-PAGE blue native polyacrylamide gel electrophoresis; GDP guanidine diphosphate; BeF_3_ berilium fluoride; RFP red fluorescent protein; CFP cyan fluorescent protein; NG mNeonGreen; MIB mitochondrial isolation buffer; NA numerical aperture;

## MATERIALS AND METHODS

### Plasmids

The following plasmids were purchased from Addgene: pBABE-hygro (#1765), pBABE-puro (#1764), pclbw-mito TagRFP (#58425), pclbw-mitoCFP (#58426). To construct the chimeric expression constructs, Mitofusin fragments were PCR amplified with overlapping sequence. The fragments were assembled into pBABE-hygro vector by SOEing PCR or Gibson assembly with 20-30 nucleotide overlap for each junction. Chimera1 is composed of Mfn1(M1-V333)+Mfn2(K355-R757); Chimera2 is composed of Mfn1(M1-E579)+Mfn2(L599-R757); Chimera3 is composed of Mfn1(M1-V333)+Mfn2(K355-C535)+Mfn1(S514-S741); Chimera4 is composed of Mfn1(M1-C513)+Mfn2(A536-T611)+Mfn1(S593-S741); Chimera5 is composed of Mfn2(M1-V354)+Mfn1(K334-C514)+Mfn2(A535-R757). Following digestion with DpnI to remove template DNA, the amplified DNA was transformed into DH5-alpha *E. coli* cells and plasmids were purified from selected colonies. Mutations in Mfn2 or Chi5 were generated in a similar approach by Gibson mutagenesis. All plasmids were confirmed by sequence analysis.

### Cell culture

All cells were grown at 37°C and 5% CO_2_ and cultured in DMEM (Thermo Fisher Scientific) containing 1X GlutaMAX (Thermo Fisher Scientific) with 10% FBS (Seradigm) or 15% FBS for Mfn1/2-null mouse embryonic fibroblasts and 1% penicillin/streptomycin (Thermo Fisher Scientific). Mouse embryonic fibroblasts cells (Mfn wild type, Mfn1-null, Mfn2-null and Mfn1/2-null) were purchased from ATCC. Cells were tested for mycoplasma contamination by DAPI staining before and after each experiment.

### Retroviral transduction and generation of clonal populations

Plat-E cells (Cell Biolabs) were maintained in complete media supplemented with 1 μg/mL puromycin and 10 μg/mL blasticidin and plated at approximately 80% confluency the day prior to transfection. Plat-E cells were transfected with FuGENE™ HD (Promega) and transfection regent was incubated overnight before a media change. Viral supernatants were collected at approximately 48, 56, 72, and 80 hours post transfection and incubated with MEFs in the presence of 8 mg/ml polybrene.

Approximately 16 hours after the last viral transduction, MEF cells were split and selection was added if needed (1 μg/mL puromycin or 200 μg/mL hygromycin).

Clonal populations were generated by plating cells at very low density and clones were collected onto sterile filter paper dots soaked in trypsin. Following expansion, whole cell extract from clonal populations were screened by western blot analysis for Mitofusin against wild type controls.

### SDS-PAGE, Western blot analysis and quantification

Following separation by SDS-PAGE, proteins transferred to nitrocellulose were detected using primary rabbit or mouse antibodies and visualized with appropriate secondary antibodies conjugated to IRDye (Thermo Fisher Scientific). Quantification was performed with Image J software (NIH).

The following antibodies were used in this study: mouse monoclonal anti-FLAG (Sigma)(1:1000); mouse monoclonal anti-Mfn2 (Sigma clone 4H8)(1:1000); mouse monoclonal anti-tubulin (Thermo Fisher Scientific clone DM1A)(1:5000); rabbit polyclonal anti-Mfn1 (gift from Jodi Nunnari)(1:500); VDAC (Thermo Fisher Scientific, polyclonal PA1-954A)(1:1000). Briefly, Mfn1 antiserum was raised against His_6_-tagged fusion proteins comprised of full-length mouse dihydrofolate reductase and an internal region of Mfn1 (residues 350-580). Fusion proteins were purified on nickel nitrilotriacetic acid columns (Thermo Fisher Scientific) in 8 M urea and eluted with 0.1% SDS and 10 mM Tris-Cl, pH 7.4.

### Transfection and microscopy

All cells were plated in No. 1.5 glass-bottomed dishes (MatTek). Mouse embryonic fibroblasts were incubated with 0.1 μg/mL Mitotracker Red CMX Ros for 15 minutes at 37°C with 5% CO_2_, washed and incubated with complete media for at least 45 minutes prior to imaging. A Z-series with a step size of 0.3 μm was collected with a Nikon Ti-E widefield microscope with a 63X NA 1.4 oil objective (Nikon), a solid-state light source (Spectra X, Lumencor), and an sCMOS camera (Zyla 5.5 Megapixel). Each cell line was imaged on at least three separate occasions (n > 100 cells per experiment).

### Image analysis

Images were deconvolved using 15 iterations of 3D Landweber deconvolution. Deconvolved images were then analyzed using Nikon Elements software. Maximum intensity projections were created using ImageJ Software (NIH). Mitochondrial morphology in mammalian cells was scored as follows: reticular indicates that fewer than 30% of the mitochondria in the cell were fragments (fragments defined as mitochondria less than 2.5 μm in length); short indicates mitochondria that were between 2.5 and 7.5 μm in length; fragmented indicates that most of the mitochondria in the cell were less than 2.5 μm in length; clustered indicates that the mitochondrial distribution was altered such that most mitochondria were coalesced in the perinuclear space.

### Preparation of mitochondria

For each experiment, three to five 15 cm plates each of MEFs stably expressing either mitochondria-targeted TagRFP or CFP were grown to `•`90% confluency. Cells were harvested by cell scraping, pelleted, and washed in mitochondrial isolation buffer (MIB) (0.2 M sucrose, 10 mM Tris-MOPS [pH 7.4], 1 mM EGTA). The cell pellet was resuspended in one cell pellet volume of cold MIB, and cells were homogenized by ∼15 strokes on ice with a Kontes Potter-Elvehjem tissue grinder set at 400 RPM. The homogenate was centrifuged (500 × *g*, 5 min, 4°C) to remove nuclei and unbroken cells, and homogenization of the pellet fraction was repeated followed by centrifugation at 500 × *g*, 5 min, 4°C. The supernatant fractions were combined and centrifuged again at 500 × *g*, 5 min, 4°C to remove remaining debris. The supernatant was transferred to a clean microfuge tube and centrifuged (7400 × *g*, 10 min, 4°C) to pellet a crude mitochondrial fraction. The crude mitochondrial pellet was resuspended in a small volume of MIB. Protein concentration of fractions was determined by Bradford assay (Bio-Rad Laboratories).

### In vitro mitochondrial fusion

An equivalent mass (12.5 - 15 μg) of TagRFP and CFP mitochondria were mixed, washed in 100 μl MIB and concentrated by centrifugation (7400 × *g*, 10 min, 4°C). Following a 10 minute incubation on ice, the supernatant was removed and the mitochondrial pellet was resuspended in 10 μl fusion buffer (20 mM PIPES-KOH [pH 6.8], 150 mM KOAc, 5 mM Mg(OAc)_2_, 0.4 M sorbitol, 0.12 mg/ml creatine phosphokinase, 40 mM creatine phosphate, 1.5 mM ATP, 1.5 mM GTP). Fusion reactions were incubated at 37°C for 60 minutes.

### Analysis of in vitro mitochondrial fusion

Mitochondria were imaged on microscope slides by mixing 3 μl fusion reaction with 3 μl of 3% low-melt agarose made in modified fusion buffer (20 mM PIPES-KOH [pH 6.8], 150 mM KOAc, 5 mM Mg(OAc)_2_, 0.4 M sorbitol) before overlaying with a coverslip. A Z-series of 0.2 μm steps was collected with a Nikon Ti-E widefield microscope with a 100X NA 1.4 oil objective (Nikon), a solid state light source (Spectra X, Lumencor), and a sCMOS camera (Zyla 5.5 Megapixel). For each condition tested, mitochondrial fusion was assessed by counting ≥ 300 total mitochondria per condition from ≥ 4 images of each condition, and individual fusion events were scored by colocalization of the red and cyan fluorophores in three dimensions.

### Protease Protection

Freshly isolated mitochondria (∼ 25 μg) were resuspended in 500 l of MIB (0.2 M sucrose, 10 mM Tris-MOPS [pH 7.4]), hypotonic mitoplast buffer (20 mM HEPES pH 7.4), or solubilizing buffer (MIB with 1% triton-X-100). After a 15 minute incubation on ice, outer mitochondrial membranes in the hypotonic buffer sample were disrupted by gently pipetting up and down 15 times. PK was added to the indicated samples to a final concentration of 100 g/mL and samples were incubated for 15 minutes on ice. PMSF was added to a final concentration of 2 mM and samples were incubated on ice for 5 minutes to stop the reaction. Mitochondria were isolated by spinning the samples at 10,400 × *g* for 15 minutes. Protein was precipitated from the solubilized sample by addition of trichloroacetic acid (TCA) to a final concentration of 12.5% v/v followed by incubation on ice for 20 minutes. The proteins were pelleted by centrifugation at 16,000 × *g* for 15 minutes, washed with ice cold acetone and dried. Mitochondrial and protein pellets were resuspended in SDS sample buffer and samples were analyzed by SDS-PAGE and Western blotting.

### Co-immunoprecipitation

Mitochondria isolated from clonal populations of Mfn1-null cells expressing either Mfn1-FLAG, Mfn2-FLAG, Chi3-FLAG or Chi5-FLAG at near endogenous levels were incubated at 37°C for 30 minutes with 2 mM GTP in Resuspension buffer (0.2 M sucrose, 20 mM HEPES-KOH [pH 7.4], 50 mM MgCl_2_). Mitochondria were then solubilized in lysis buffer (20 mM HEPES-KOH [pH 7.4], 50 mM KCl, 5 mM MgCl_2_) with 1.5% n-Dodecyl β-D-maltoside (DDM) and 1X Halt Protease Inhibitor (Thermo Scientific) for 30 minutes on ice. Lysates were cleared at 10,000 × g for 15 minutes at 4°C. Supernatant was incubated with 50 L magnetic mMACS anti-DYKDDDDK Microbeads (Miltenyi Biotec) for 30 minutes on ice. The sample was applied to a MACS column (Miltenyi Biotec) and washed twice with Wash I (20 mM HEPES-KOH [pH 7.4], 50 mM KOAc, 5 mM MgCl_2_, 0.1% DDM) and once with Wash II (20 mM HEPES-KOH [pH 7.4], 50 mM KCl, 5 mM MgCl_2_). One column volume (25 μl) SDS-PAGE loading buffer (60 mM Tris-HCl [pH 6.8], 2.5% sodium dodecyl sulfate, 5% βME, 5% sucrose, 0.1% bromophenol blue) was incubated for 15 minutes at room temperature and proteins were eluted twice with 50 μL SDS-PAGE loading buffer. The majority of the protein eluted in the first 50 μL elution. Samples were run on an SDS-PAGE gel and transferred onto nitrocellulose. Membranes were blocked in 4% Milk for at least 45 minutes and were probed with anti-Mfn2, anti-FLAG or anti-VDAC antibody for 4 hours at room temperature or overnight at 4°C. Membranes were incubated with DyLight secondary antibody (Invitrogen) at room temperature for 45 minutes. Membranes were imaged on LI-COR Imaging System (LI-COR Biosciences).

### BN-PAGE

Isolated mitochondria (10 – 30 μg) were incubated with or without 2 mM nucleotide and 2.5 mM BeSO_4_, and 25 mM NaF as indicated at 37°C for 30 minutes. Mitochondria were then lysed in 1% w/v digitonin, 50 mM Bis-Tris, 50 mM NaCl, 10% w/v glycerol, 0.001% Ponceau S; pH 7.2 for 15 minutes on ice. Lysates were centrifuged at 16,000 × *g* at 4°C for 30 minutes. The cleared lysate was mixed with Invitrogen NativePAGE^TM^ 5% G-250 Sample Additive to a final concentration of 0.25%. Samples were separated on a Novex™ NativePAGE™ 4 - 16% Bis-Tris Protein Gels (Invitrogen) at 4°C. Gels were run at 40 volts for 30 minutes then 100 volts for 30 minutes with dark cathode buffer (1X NativePAGE^TM^ Running Buffer [Invitrogen], 0.02% [w/v] Coomassie G-250). Dark cathode buffer was replaced with light cathode buffer (1X NativePAGE^TM^ Running Buffer [Invitrogen], 0.002% [w/v] Coomassie G-250) and the gel was run at 100 volts for 30 minutes and subsequently at 250 volts for 60-75 minutes until the dye front ran off the gel. After electrophoresis was complete, gels were transferred to PVDF membrane (Bio-Rad Laboratories) at 30 volts for 16 hours in transfer buffer (25 mM Tris, 192 mM glycine, 20% methanol). Membranes were incubated with 8% acetic acid for 15 minutes and washed with H_2_O for 5 minutes. Membranes were dried at 37°C for 20 minutes and then rehydrated in 100% methanol and washed in H_2_O. Membranes were blocked in 4% milk for 20 minutes and were probed with anti-FLAG (Sigma) for 4 hours at room temperature or overnight at 4°C. Membranes were incubated with HRP-linked secondary antibody (Cell Signaling Technology) at room temperature for 1 hour. Membranes were developed in SuperSignal Femto ECL reagent (Thermo Fisher Scientific) for 5 minutes and imaged on iBright Imaging System (Thermo Fisher Scientific). Band intensities were quantified using ImageJ software (NIH). NativeMark Unstained Protein Standard (Life Technologies) was used to estimate molecular weights of Mitofusin protein complexes.

## Supporting information

Sloat Supplement

## ACKNOWLEDGEMENTS

We would like to thank members of the Hoppins lab, Alex Merz, Laura Lackner and Marijn Ford for scientific suggestions and critical reading of the manuscript.

