## Supplementary material for "Identification of a Mitofusin specificity region that confers unique activities to Mfn1 and Mfn2": Sloat Supplement

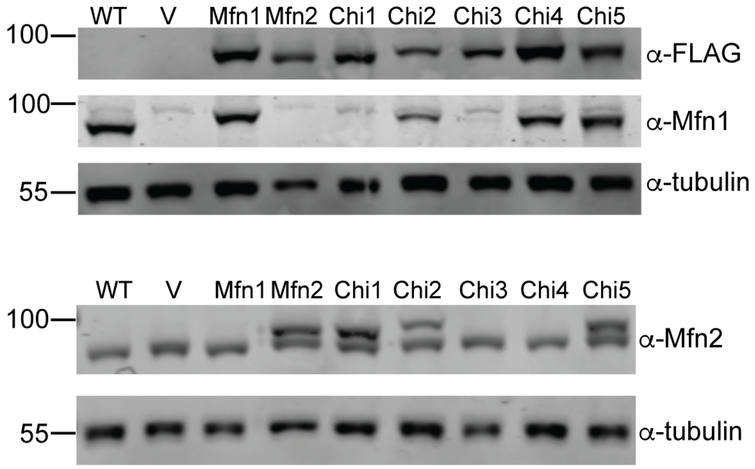


**Figure S1**

**Mitofusin-FLAG protein expression in clonal populations of Mfn1-null cells.**
Whole cell lysates prepared from the indicated cell lines were subjected to SDS-PAGE and immunoblotting with the indicated antibody. The Mfn1 epitope is amino acids 350-580 and the Mfn2 epitope is amino acids 661-757.

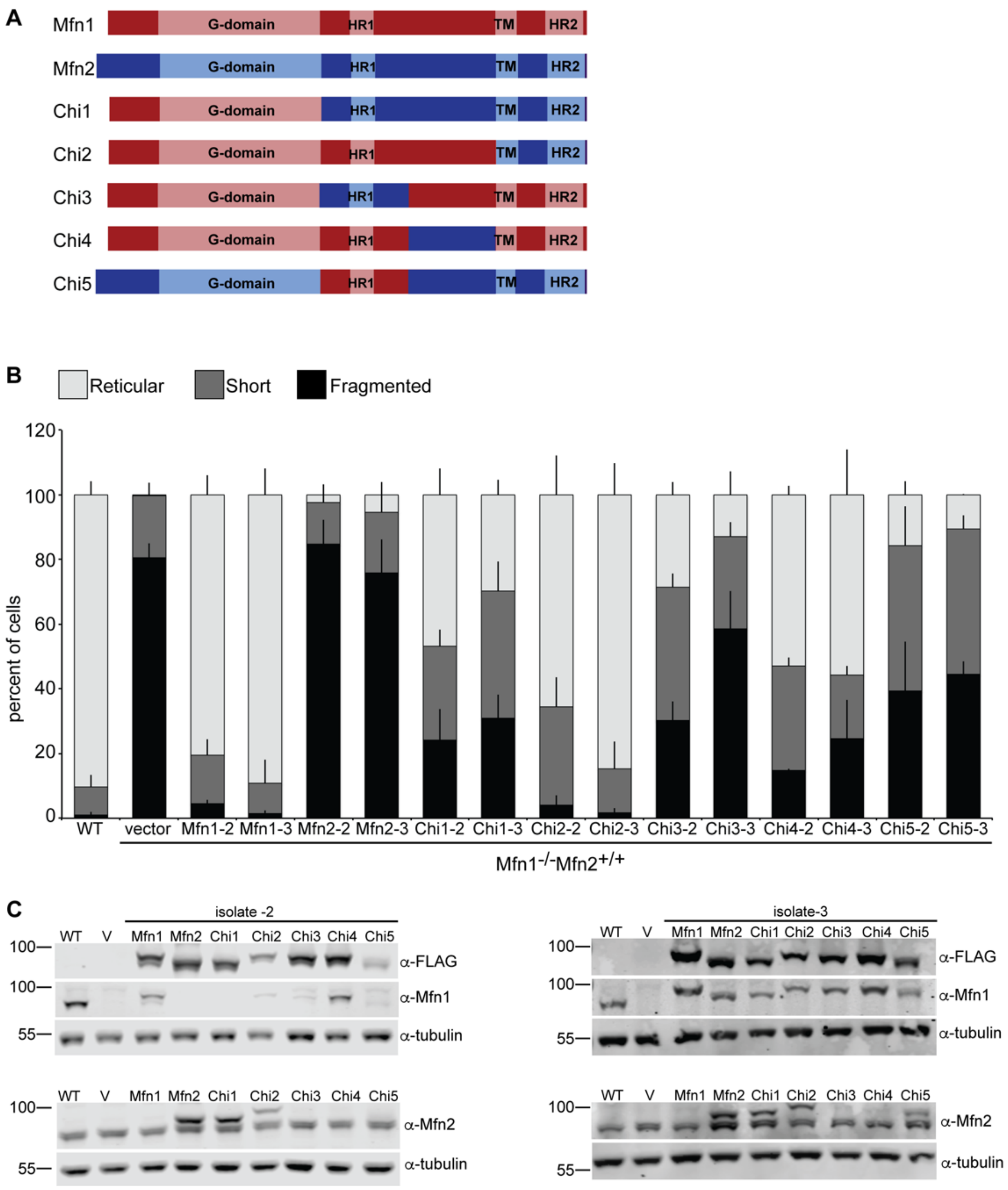


### Figure S2

**Mfn1-dependent rescue of Mfn1-null cells by Chimera proteins.** (A) Schematic representation of known functional domains in Mfn1 and Mfn2 and the chimeric proteins generated for this study. (B) Quantification of the mitochondrial morphology in a clonal population of Mfn1-null cells expressing the indicated Mitofusin or chimeric protein (-2 and -3 indicate the second and third clonal populations, respectively). Error bars indicate mean + standard deviation from three blinded experiments (n ≥ 100 cells per population per experiment). (C) Whole cell lysates prepared from the indicated cell lines described in (B) were subjected to SDS-PAGE and immunoblotting with the indicated antibody.

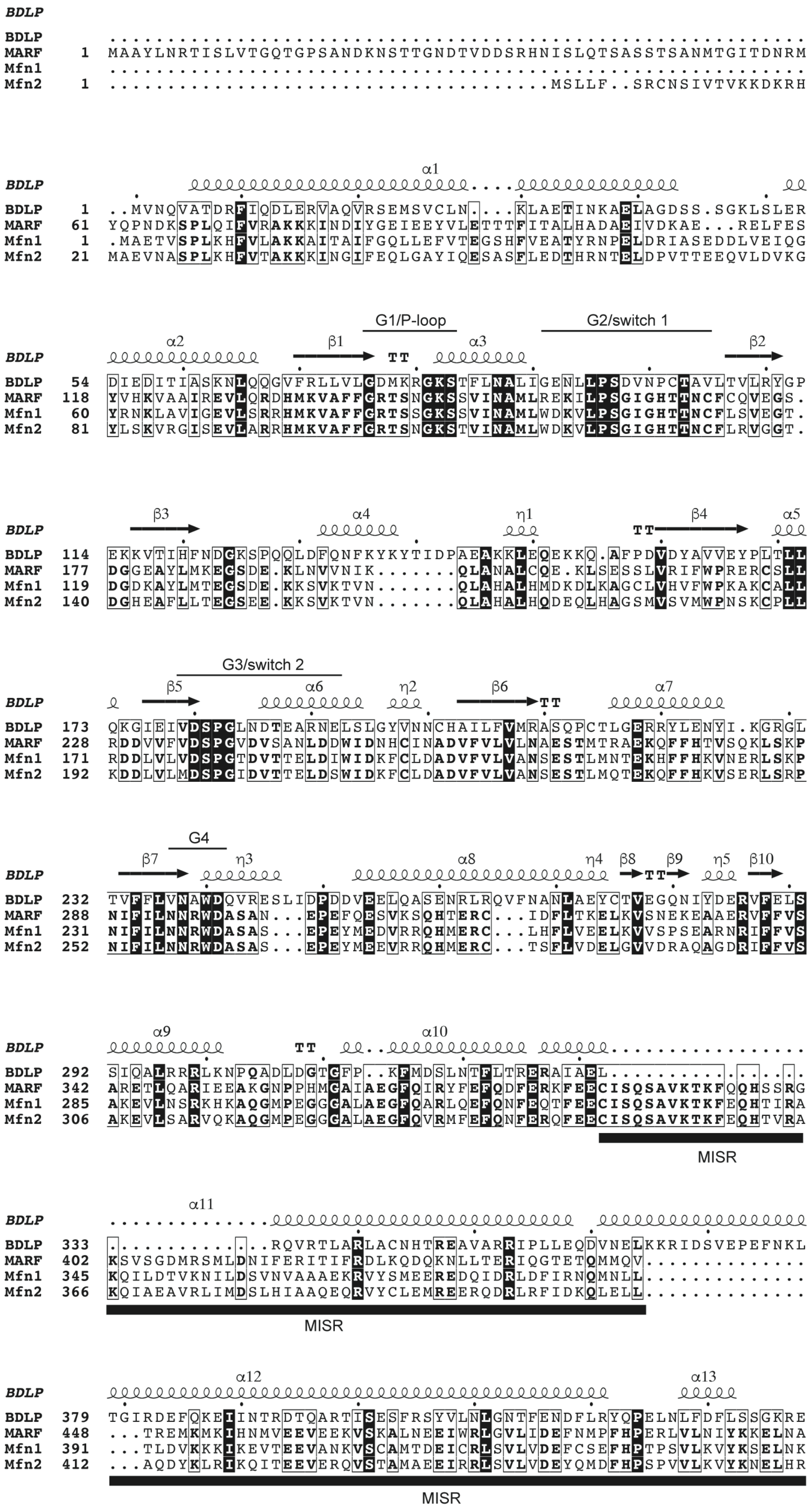
BDLP
BDLP ............................................................
MARF 1 MAAYLNRTISLVTGQTGPSANDKNSTTGNDTVDDSRHNISLQTSASSTSANMTGITDNRM
Mfn1 ............................................................
Mfn2 1 ..................................MSLLF..SRCNSIVTVKKDKRH
α1
BDLP 1 ..MVNQVATDRFIQDLERVAQVRSEMSVCLN....KLAETINKAELAGDSS.SGKLSLER
MARF 61 YQPNDKSPLQIFVRAKKKINDIYGEIEEYVLETTTFITALHADAEIVDKAE...RELFES
Mfn1 1 .MAETVSPLKHFVLAKKAITAIFGQLLEFVTEGSHFVEATYRNPELDRIASEDDLVEIQG
Mfn2 21 MAEVNASPLKHFVTAKKKINGIFEQLGAYIQESASFLEDTHRNTELDPVTTEEQVLDVKG
G1/P-loop
G2/switch 1
α2 β1 TT α3 β2
BDLP 54 DIEDITIASKNLQQGVFRLLVLGDMKRGKSTFLNALIGENLLPSDVNPCTAVLTVLRYGP
MARF 118 YVHKVAAIREVLQRDHMKVAFFGRTSNGKSSVINAMLREKILPSGIGHTTNCFCQVEGS.
Mfn1 60 YRNKLAVIGEVLSRRHMKVAFFGRTSSGKSSVINAMLWDKVLPSGIGHTTNCFLSVEGT.
Mfn2 81 YLSKVRGISEVLARRHMKVAFFGRTSNGKSTVINAMLWDKVLPSGIGHTTNCFLRVGGT.
β3 α4 η1 TT β4 α5
BDLP 114 EKKVTIHFNDGKSPQQLDFQNFKYKYTIDPAEAKKLEQEKKQ.AFPDVDYAVVEYPLTLL
MARF 177 DGGEAYLMKEGSDE.KLNVVNIK.......QLANALCQE.KLSESSLVRIFWPRERCSLL
Mfn1 119 DGDKAYLMTEGSDE.KKSVKTVN.......QLAHALHMDKDLKAGCLVHVFWPKAKCALL
Mfn2 140 DGHEAFLLTEGSEE.KKSVKTVN.......QLAHALHQDEQLHAGSMVSVMWPNSKCPLL
G3/switch 2
β5 α6 η2 β6 TT α7
BDLP 173 QKGIEIVDSPGLNDTEARNELSLGYVNNCHAILFVMRASQPCTLGERRYLENYI.KGRGL
MARF 228 RDDVVFVDSPGVDVSANLDDWIDNHCINADVFVLVLNAESTMTRAEKQFFHTVSQKLSKP
Mfn1 171 RDDLVLVDSPGTDVTTELDIWIDKFCLDADVFVLVANSESTLMNTEKHFFHKVNERLSKP
Mfn2 192 KDDLVLMDSPGIDVTTELDSWIDKFCLDADVFVLVANSESTLMQTEKQFFHKVSERLSRP
G4
β7 η3 α8 η4 β8 TT β9 η5 β10
BDLP 232 TVFFLVNAWDQVRESLIDPDDVEELQASENRLRQVFNANLAEYCTVEGQNIYDERVFELS
MARF 288 NIFILNNRWDASAN...EPEFQESVKSQHTERC...IDFLTKELKVSNEKEAAERVFFVS
Mfn1 231 NIFILNNRWDASAS...EPEYMEDVRRQHMERC...LHFLVEELKVVSPSEARNRIFFVS
Mfn2 252 NIFILNNRWDASAS...EPEYMEEVRRQHMERC...TSFLVDELGVVDRAQAGDRIFFVS
α9 TT α10
BDLP 292 SIQALRRLKNPQADLDGTGFP..KFMDSLNTFLTRERAIAEL..................
MARF 342 ARETLQARIEEAKGNPPHMGAIAEGFQIRYFEFQDFERKFEECISQSAVKTKFQQHSSRG
Mfn1 285 AKEVLNSRKHKAQGMPEGGGALAEGFQARLQEFQNFEQTFEECISQSAVKTKFEQHTIRA
Mfn2 306 AKEVLSARVQKAQGMPEGGGALAEGFQVRMFEFQNFERQFEECISQSAVKTKFEQHTVRA
MISR
α11
BDLP 333 ..............RQVRTLARLACNHTREAVARRIPLLEQDVNELKKRIDSVEPEFNKL
MARF 402 KSVSGDMRSMLDNIFERITIFRDLKQDQKNLLTERIQGTETQMMQV..............
Mfn1 345 KQILDTVKNILDSVNVAAAEKRVYSMEEREDQIDRLDFIRNQMNLL..............
Mfn2 366 KQIAEAVRLIMDSLHIAAQEQRVYCLEMREERQDRLRFIDKQLELL..............
MISR
α12 α13
BDLP 379 TGIRDEFQKEIINTRDTQARTISESFRSYVLNLGNTFENDFLRYQPELNLFDFLSSGKRE
MARF 448 ...TREMKMKIHNMVEEVEEKVSKALNEEIWRLGVLIDEFNMPFHPERLVLNIYKKELNA
Mfn1 391 ...TLDVKKKIKEVTEEVANKVSCAMTDEICRLSVLVDEFCSEFHPTPSVLKVYKSELNK
Mfn2 412 ...AQDYKLRIKQITEEVERQVSTAMAEEIRRLSVLVDEYQMDFHPSPVVLKVYKNELHR
MISR

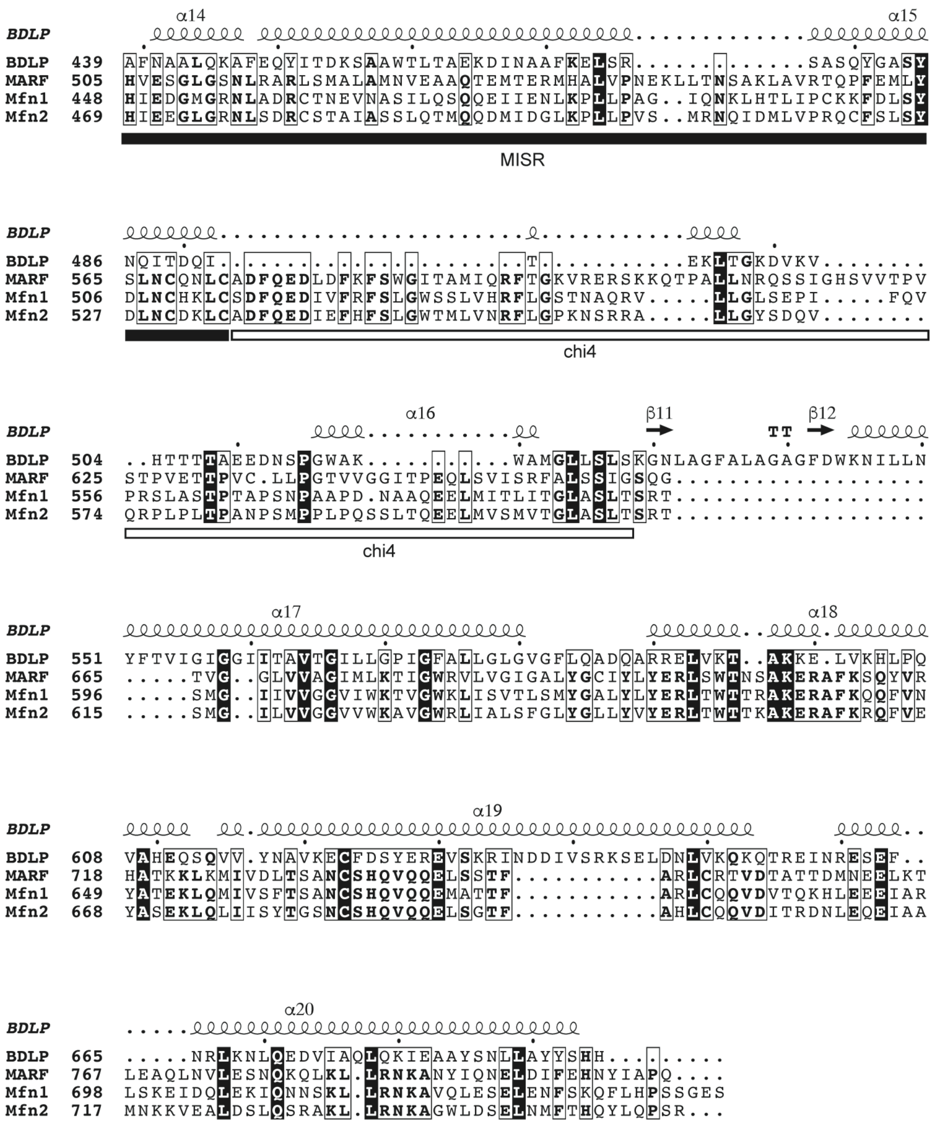

**Figure S3**

**Sequence alignment of BDLP, MARF and the mouse Mitofusin proteins.** Amino acid sequences of BDLP (B2IZD3), MARF (Q7YU24), Mfn1 (Q811U4) and Mfn2 (Q80U63) were aligned with Clustal Omega. The structural features of BDLP PDB 2J68 are shown: alpha helices are indicated with spirals and beta sheets with arrows. The graphic was generated by ESPript (Robert and Gouet, 2014). The G1-G4 elements are specified above the sequence. MISR and the Chi5 regions are identified below the sequence.

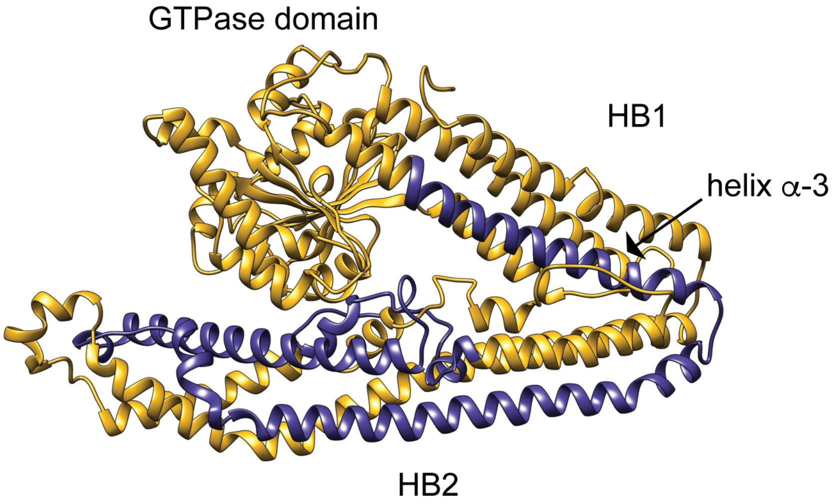


**Figure S4**

**BDLP structure as a model for Mitofusin structure.**

A homology model of Mfn1 generated by Phyre2 utilizing PDB 2J68 (Kelley et al., 2015). MISR is highlighted in purple and includes helix $\alpha$-3 in HB1 of the internally truncated Mfn1 structure (Cao et al., 2017, Qi et al., 2016, Yan et al., 2018)as well as predicted helices and loops in the uncharacterized HB2.

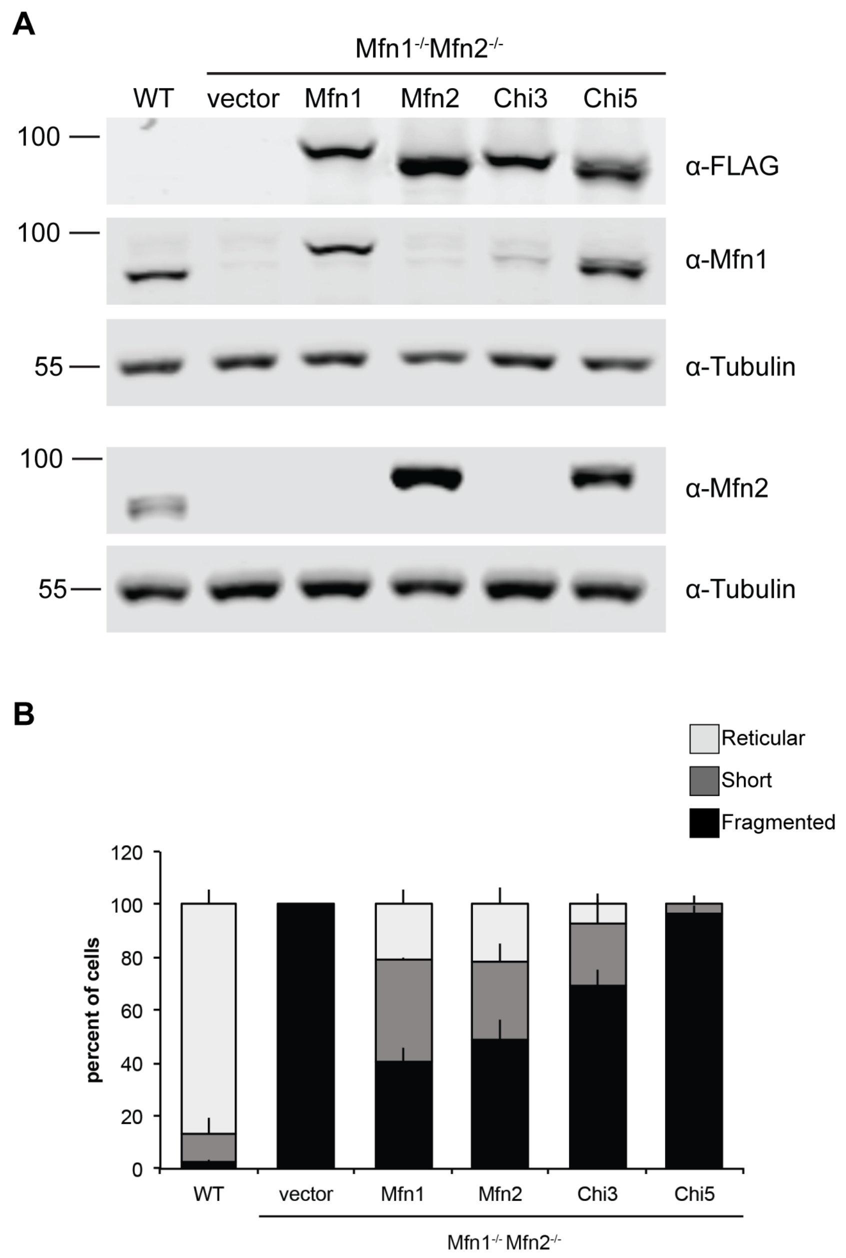


**Figure S5**

**Mitofusin-FLAG protein expression in clonal populations of Mfn1/2-null cells.**

(A) Whole cell lysates prepared from the indicated cell lines were subjected to SDS-PAGE and immunoblotting with the indicated antibody. (B) Quantification of the mitochondrial morphology in a clonal population of Mfn1/2-null cells expressing the indicated Mitofusin or Chimeric protein.

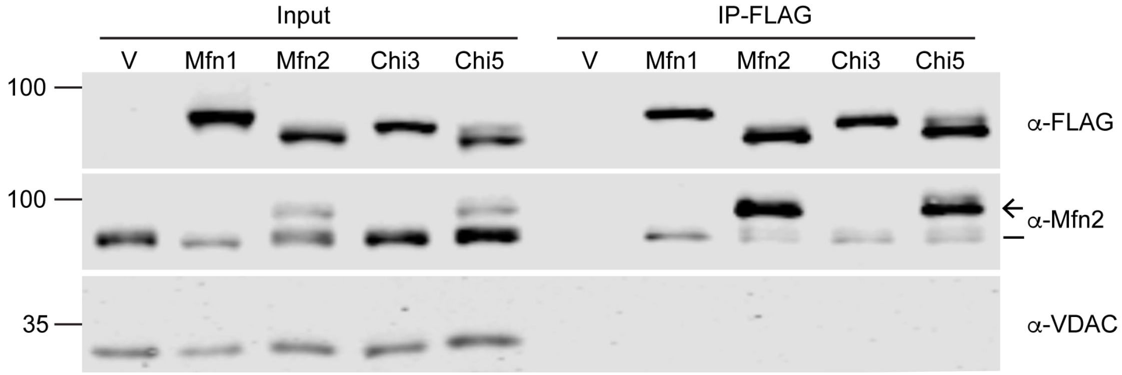


**Figure S6**

**Co-immunoprecipitation of Mitofusin-FLAG and endogenous Mfn2.**

Immunoprecipitations were performed with anti-FLAG magnetic beads on 100 μg of mitochondria isolated from Mfn1-null cells expressing the indicated Mitofusin-FLAG. Samples were analyzed by SDS-PAGE and Western blot with the indicated antibodies. For α-Mfn2, the line indicates the endogenous protein and the arrow indicates the FLAG-tagged variant. Input represents 2.5% of the total lysate and IP-FLAG represents 50% of the immunoprecipitated protein.

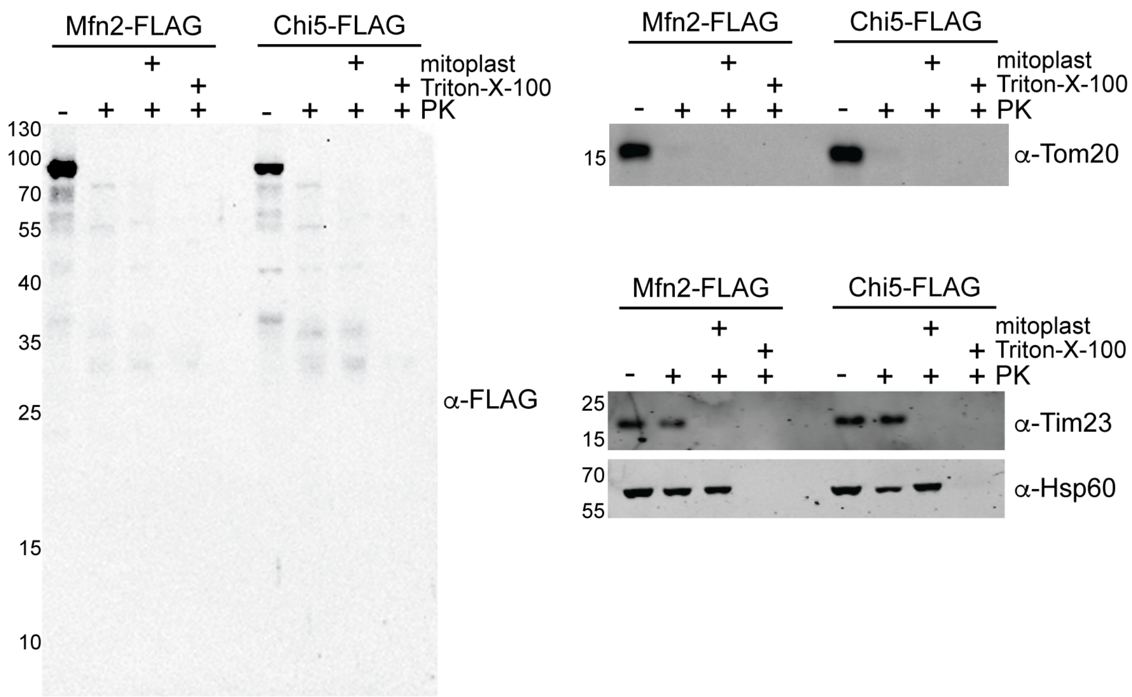


**Figure S7**

**Protease protection of Mfn2 and Chi5**.

Mitochondria were isolated from Mfn1-null cells expressing either Mfn2-FLAG or Chi5-FLAG, as described in Fig. 1 and Fig. S1. Mitochondria were left intact, converted to mitoplasts or solubilized with Triton-X-100 before treatment with (+) or without proteinase K (PK). Following treatment, samples were analyzed by SDS-PAGE and Western blot analysis against the indicated proteins. Tom20 represents a mitochondrial outer membrane marker; Tim23 represents a mitochondrial inner membrane marker; Hsp60 represents a mitochondrial matrix marker.
